# 17-(Allylamino)-17demethoxygeldanamycin reduces Endoplasmic Reticulum (ER) stress-induced mitochondrial dysfunction in C2C12 myotubes

**DOI:** 10.1101/350702

**Authors:** Adam P. Lightfoot, Rhiannon S. Morgan, Joanna E. Parkes, Anastasia Thoma, Lesley A. Iwanejko, Robert G. Cooper

## Abstract

In patients with myositis, persistent skeletal muscle weakness in the absence of significant inflammatory cell infiltrates is a well-recognised, but poorly understood, cause of morbidity. This has led researchers to investigate cellular mechanisms independent of immune cells, which may contribute to this underlying muscle weakness. Chronic ER stress pathway activation is evident in the muscle of myositis patients, and is now a potential mediator of muscle weakness in the absence of inflammation. Abnormal ER stress pathway activation is associated with mitochondrial dysfunction, resulting in bioenergetic deficits and reactive oxygen species (ROS) generation, which in this context may potentially damage muscle proteins and thus impair contractile performance. This study examined whether treatment with the HSP90 inhibitor 17-*N*-allylamino-17-demethoxygeldanamycin (17AAG) could mitigate these ER stress-induced changes. C2C12 myotubes were treated with the ER stress-inducing compound Tunicamycin, in the presence or absence of 17AAG. Myotubes were examined for changes relating to ER stress pathway activation, mitochondrial function, markers of oxidative damage and in myotubular dimensions. ER stress pathway activation caused mitochondrial dysfunction, as evidenced by reduced oxygen consumption and ATP generation and by increased gene expression levels of the bio-energetic regulator, uncoupling protein 3 (*UCP-3)*, the latter indicative of electron transport chain uncoupling. ER stress pathway activation also caused increased gene expression of superoxide dismutase (*SOD) 2* and peroxiredoxin (*PRDX) 3*, elevated H_2_O_2_ levels, and reduced total thiol pool levels and a significant diminution of myotubular dimensions. Exposure to 17AAG ameliorated these ER stress-induced changes. These findings, which suggest that 17AAG can reduce ER stress-induced mitochondrial dysfunction, oxidative damage and myotubular atrophy, have potential implications in the context of human myositis.

## Background

The idiopathic inflammatory myopathies, collectively termed myositis are characterised by infiltrations of T and B cells preferentially into the proximal muscles, to cause myofibre damage and debilitating proximal muscle weakness. A histological hallmark of myositis is the detection of up-regulated human leukocyte antigen (HLA) I, both within and on the cell surface of muscle fibres seen in diagnostic muscle biopsies, [1]. Traditionally it has been assumed that infiltration of inflammatory T and B cells in myositis represents the key pathological mechanism responsible for damaging myofibres, and infiltrating inflammatory cells clearly do have this potential, [2]. However, that inflammatory cells are the sole pathogenic effector causing myofibre damage is being challenged by accumulating evidence that non-immune cell-mediated factors are also implicated, [3].

The inflammatory cell loads detected in diagnostic muscle biopsies often correlate only poorly with the severity of myositis patients’ demonstrable weakness, while weakness frequently persists in patients with apparently well suppressed muscle inflammation, [4] [5]. Furthermore, in immune-mediated necrotising myopathy, a myositis subtype so called because it associates strongly with the presence of anti-SRP or anti-HMGCoA reductase autoantibodies and marked creatine kinase elevations, severe myonecrosis occurs in the absence or marked paucity of infiltrating B and T cells, [6]. Lastly, in a murine model of inflammatory myositis specifically induced by transgenic up-regulation of HLA I, weakness may actually precede inflammatory cell infiltrations, [4]. These findings clearly suggest that myotoxic factors other than immune cell infiltrates are also pathologically involved. Recent reviews have thus suggested that non-immune cell-mediated mechanisms must also play a significant cytotoxic role, over and above that expected from infiltrating inflammatory cells, [7, 8]. Research in human and murine myositis models suggests that chronic over-activation of the ER stress pathway also contributes to weakness induction, [9]. However, the precise mechanisms which mediate weakness induction downstream of ER stress pathway activation remain poorly understood, [7].

Studies in muscle and non-muscle cells demonstrate that ER stress pathway activation directly influences mitochondrial function, [10]. Pathway activation physiologically causes calcium ion release, which, when taken up by adjacent mitochondria, causes bioenergetic changes, including changes in ROS generation and ATP synthesis rates, [10–12]. These are normal functions, but when ER pathway activation is abnormal and/or prolonged, as occurs in myositis, growing evidence suggests that this may adversely affect mitochondrial function, [13, 14]. Research from this and other laboratories has demonstrated that targeted up-regulation of cytoprotective chaperones, i.e. heat shock proteins (HSPs), can potentially alleviate oxidative damage to skeletal muscle fibres from otherwise unchecked ROS generation, [15, 16]. Moreover, pharmacological up-regulation of molecular chaperones by use of 17AAG prevents contraction-induced muscle fibre damage in older mice, [17].

Given the mutual proximity of the ER and mitochondria within skeletal muscle cells, and the chronic nature of ER stress pathway activation in human myositis, we have in this study used an *in vitro* murine muscle cell model to mimic the ER stress component probably present in human myositis. The overall aim of the study was to test whether 17AAG could prevent or reduce ER stress-related mitochondrial dysfunction.

## Methods

### Chemicals and reagents

All chemicals and reagents were supplied by Sigma Aldrich, UK, unless stated otherwise.

### Cell culture, treatments, and preparation

Murine C2C12 myotubes were cultured in standard conditions (5% CO_2_, 37°C) in Dulbecco’s modified eagles medium (DMEM) supplemented with 10% foetal bovine serum (FBS) (v/v), 2mM L-glutamine and 50-i.u. penicillin. Cells were cultured to ~ 60-70% confluence, when the growth media were replaced by differentiating media, containing 2% horse serum (HS). Cells were set to differentiate into myotubes over a seven-day period, all “treatments” being deployed on day seven. At this point, myotubes were treated with Tunicamycin (0.1μg/ml) in the absence or presence of 17AAG (0.1μg/ml) for a period of 24 hours. Tunicamycin induces ER stress, by inhibiting the *n*-glycosylation step of protein folding, and resulting in misfolded protein accumulations within the ER lumen. After 24 hours of these treatments, or without either in the case of control cells, the muscle cells were harvested, using ice-cold phosphate buffered saline (PBS), and stored at −80°C until further analysis.

### Microscopy

Without and following Tunicamycin, myotubes were imaged using confocal microscopy (x10 magnification, Nikon instruments). Images were randomised and assigned to a blinded researcher for dimensional analysis. Myotubular diameters were determined using ImageJ software, measuring ten myotubes per dish. Dimensional data of treated cells were expressed as a percentage of those of the control cells, i.e. those receiving no Tunicamycin or 17AAG treatment.

### SDS-PAGE and western blotting

Proteins were extracted from myotube lysates by brief sonication in RadioImmunoPrecipitation assay (RIP A) buffer, comprising: 150mM NaCl, 0.1% Triton X-100, 0.5% deoxycholic acid, 0.1% SDS, 50mM Tris-HCl pH 8.0, supplemented with EDTA-free protease inhibitor cocktail and PhosSTOP phosphatase inhibitor cocktail, as per the manufacturer’s guidelines (Roche Pharmaceuticals). Quantification of total cellular protein was determined using the Bradford Assay, [18]. Fifty micrograms of sample was separated on 4%/12% acrylamide gels (National Diagnostics) and proteins transferred to a Polyvinylidene fluoride (PVDF) membrane by western blotting, using semi-dry transfer (Geneflow, UK). PVDF membranes were analysed using primary antibodies specific to Grp94 (1:2000), Grp78 (1:1000), IκBCα (1:1000), Total OXPHOS antibody cocktail (1:1000), Caspase-12 (1:1000), Vinculin (1:5000), beta actin (1:5000), and phosphorylated JNK (1:500), with species-specific HRP-conjugate secondary antibodies (1:5000). Enhanced chemiluminesence (ECL) (Amersham, Cardiff, UK) was used to detect bands using a Bio-Rad Chemi-doc XRS system with Imagelab software (Bio-Rad, Hercules, USA).

### qPCR

The TRIzol phenol/chloroform method was used for RNA extraction, with purification by GeneJet clean-up kit (Thermo Fisher Scientific). Complimentary DNA was synthesised using Bio-Rad iScript first strand kit, according to the manufacturer’s protocol (Bio-Rad, Heracles). Quantitative polymerase chain reaction (qPCR) was carried out using Sybr green master mix (Roche Diagnostics), in accordance with the manufacturer’s protocol; and analysis undertaken using the 2^−ΔΔct^ method, [19]. The primers used are listed in **Supplementary Table 1,** the choice of housekeeping genes (s29, RPL7, RPL32) was determined based on stability of expression in C2C12 cells spanning treatment groups.

### Mitochondrial function assays

Oxygen consumption was assessed using an Oxytherm clark electrode (Hansatech Instruments). Myotubes were harvested and suspended within the electrode chamber in *buffer z* comprising (in mM): 110 K-MES, 35 KCl, 1 EGTA, 5 K_2_HPO_4_, and 3 MgCl_2_.6 H_2_O, 0.03% fatty acid free bovine serum albumin (BSA), and 0.5μg/ml saponin, pH 7.1 at 37°C, [20]. Maximal respiration was induced by introducing 15mM of ADP, representing state 3 respiration; following depletion of ADP, cells revert to state 4 respiration, [21]. The respiratory control ratio (RCR) was determined by dividing the active respiration rate (state III) by the rate following ADP depletion (state IV). Note state IV respiration is not equivalent to basal respiration, typically occurring at a faster rate due to endogenous ATPase activity in cells, breaking down ATP generated to ADP (**Figure 1**). The phosphate:oxygen (P:O) ratio is the relationship between oxygen consumption and ATP synthesis, and was calculated by determining the volume of oxygen consumed during maximal respiration, [21]. Respirometry data were normalised to total cellular protein levels, the latter as determined by the Bradford Assay. Total myotubular ATP levels were determined using an ATP Bioluminescence assay kit HS II (Roche Pharmaceuticals), in accordance with the manufacturer’s protocol. Total ATP concentrations were normalised to total cellular protein levels, as per the Bradford assay. Total protein thiols (sulphdryl) were quantified in the myotube harvests in 1% 5-sulfosalicylic acid (SSA) solution, and re-suspended in Tris/HCl buffer, as described by Di Monte et al, [22]. Total thiol levels were normalised to total cellular protein levels, as per the Bradford assay.

**Figure 1:**
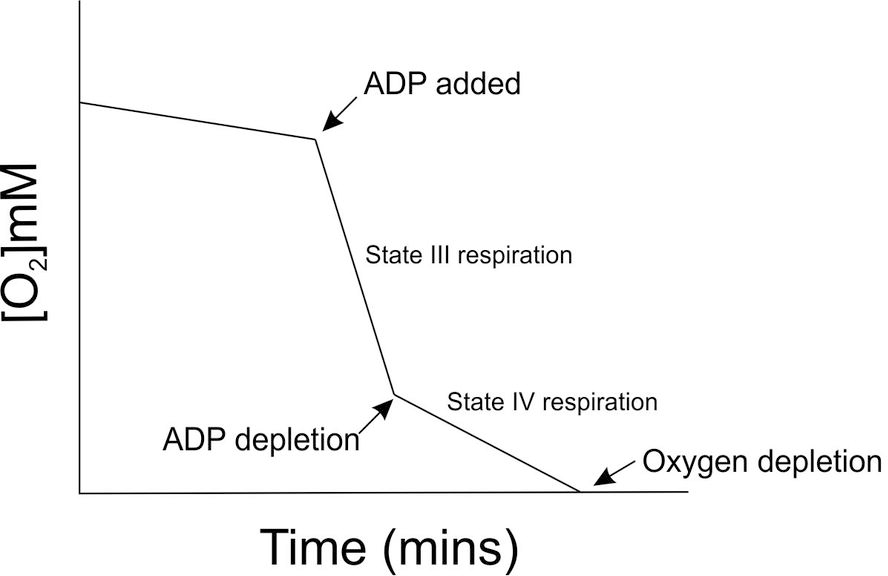
Schematic illustrating mitochondrial respiration states [O_2_], 15mM ADP was added to induce active respiration (state 3), and tracked over time through the depletion of ADP, driving the cells into state 4 respiration and finally oxygen depletion in the system. Values derived from the oxygraph experiment were used for the calculation of RCR and P:O.

### Amplex Red assay

Hydrogen peroxide levels in C2C12 myotubes treated with Tunicamycin, with or without 17AAG were quantified by the Amplex Red assay (Thermo Fisher Scientific), in accordance with the manufacturer’s protocol, [23].

### Phosphorylated JNK ELISA

Following Tunicamycin treatment, without or with 17AAG, whole cell lysates were isolated from the myotubes, and ELISA (eBioscience) used to detect phosphorylated levels of Jun N-terminal kinase (JNK), in accordance with the manufacturer’s protocol. Data were normalised to total cellular protein levels, as per the Bradford assay.

### Statistical Analysis

Data are presented as mean ± SEM; statistical analyses made using analysis of variance (ANOVA), followed by *post hoc* least significant difference testing or Kruskal-Wallis test where appropriate. Data were analysed using SPSS software, and p values ≤ 0.05 considered as statistically significant.

## Results

### 17AAG protects against ER stress-induced myotubular atrophy, but does not alter Tunicamycin-induced ER stress pathway activation

Doses of 1.0 and 10 μg/ml of Tunicamycin induced significant myotubular caspase-12 cleavage (**Figure 2A**); indicating that apoptosis had been induced at these dosages. A Tunicamycin dose of 0.1μg/ml did not induce significant caspase-12 cleavage, confirming that apoptosis had not occurred. Thus, the term Tunicamycin hereafter always refers to this non-apoptotic dose (0.1μg/ml). Treatment of myotubes with this Tunicamycin dose induced significant increases in protein levels of Grp78 and Grp94 (**Figure 2B-C**), and up-regulation of gene levels of spliced *XBP-1* (**Figure 2C**), results clearly confirming that ER stress had been induced, [24]. Grp94 protein content was significantly reduced when tuniamcyin was incubated in the presence of 17AAG (**Figures 2B**).

**Figure 2:**
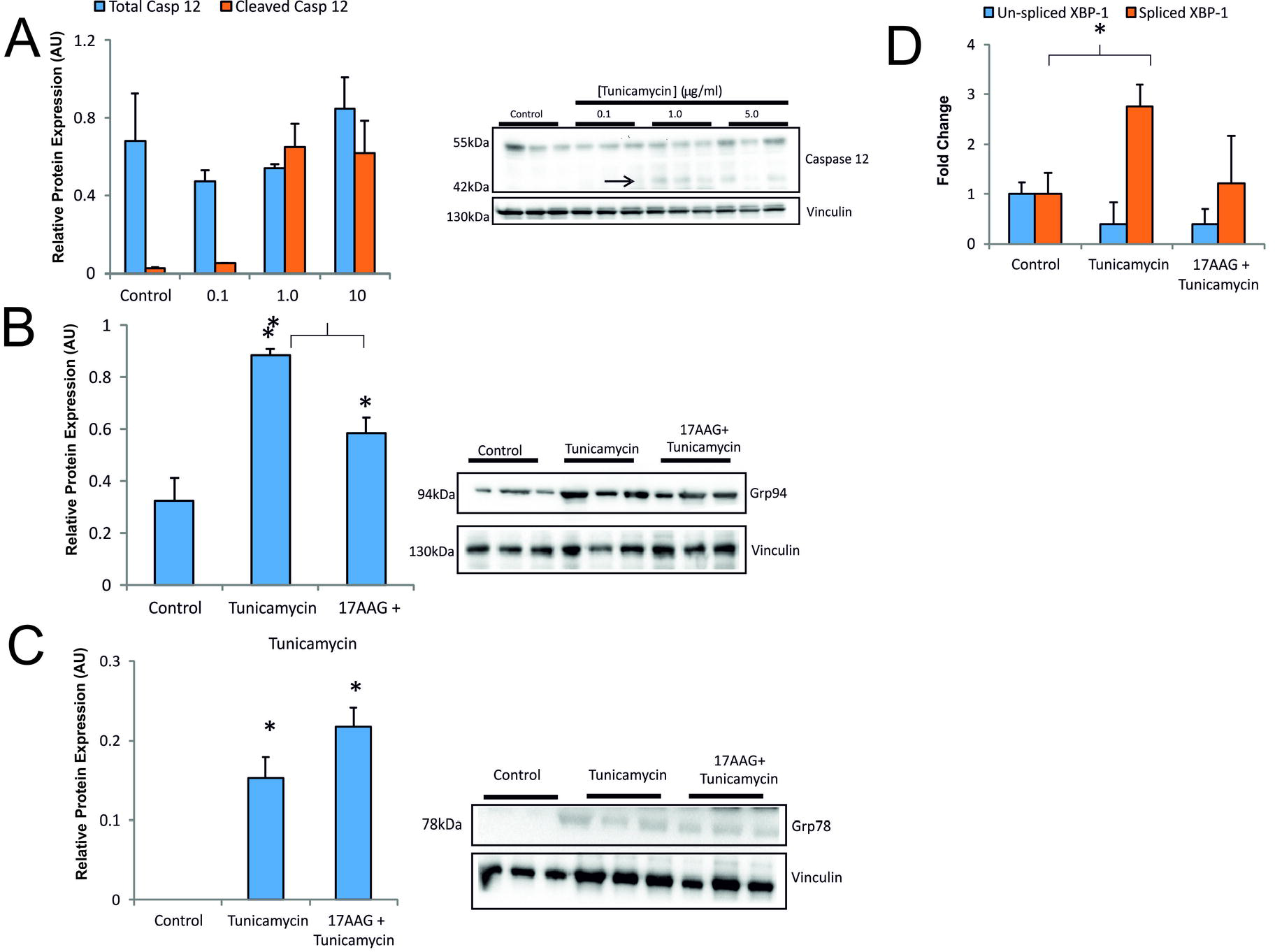
Representative western blot images and quantified densitometry showing levels from myotubes treated with Tunicamycin, without and with 17AAG. Results of: **(A)** Caspase-12 (full and cleaved forms) – The increases in the cleaved forms at the 1.0 and 10 μg/ml doses confirmed that apoptosis had been induced at these dosages, while the unchanged cleaved form levels at the 0.1ug/ml dose confirmed this as a non-apoptotic dosage; **(B)** Grp94 – The increases with Tunicamycin confirmed that ER stress had been induced **(C)** Grp78 – The increases with Tunicamycin confirmed that ER stress had been induced, but unaffected by 17AAG; **(D)**; qPCR (un-spliced and spliced *XBP-1* gene expression) – The increases with Tunicamycin confirmed that ER stress had been induced. Data presented are mean ± SEM (n=3-6) *p≤0.05.

Treatment with non-apoptotic Tunicamcyin doses also induced significant reductions in myotubular diameters (**Figures 3A-B**). Given these dimensional reductions, we interrogated for the mechanisms responsible. Surprisingly, no changes were detected in the gene expression levels of the ubiquitin ligases *Atrogin-1* and *MuRF-1*, both of which are known to mediate muscle fibre atrophy, [25] (**Figure 3C**). In contrast, Tunicamycin induced a significant reduction in I*κ*Bα levels (see later), an effect reduced when Tunicamcyin was given with 17AAG (**Figures 3D-E**).

**Figure 3:**
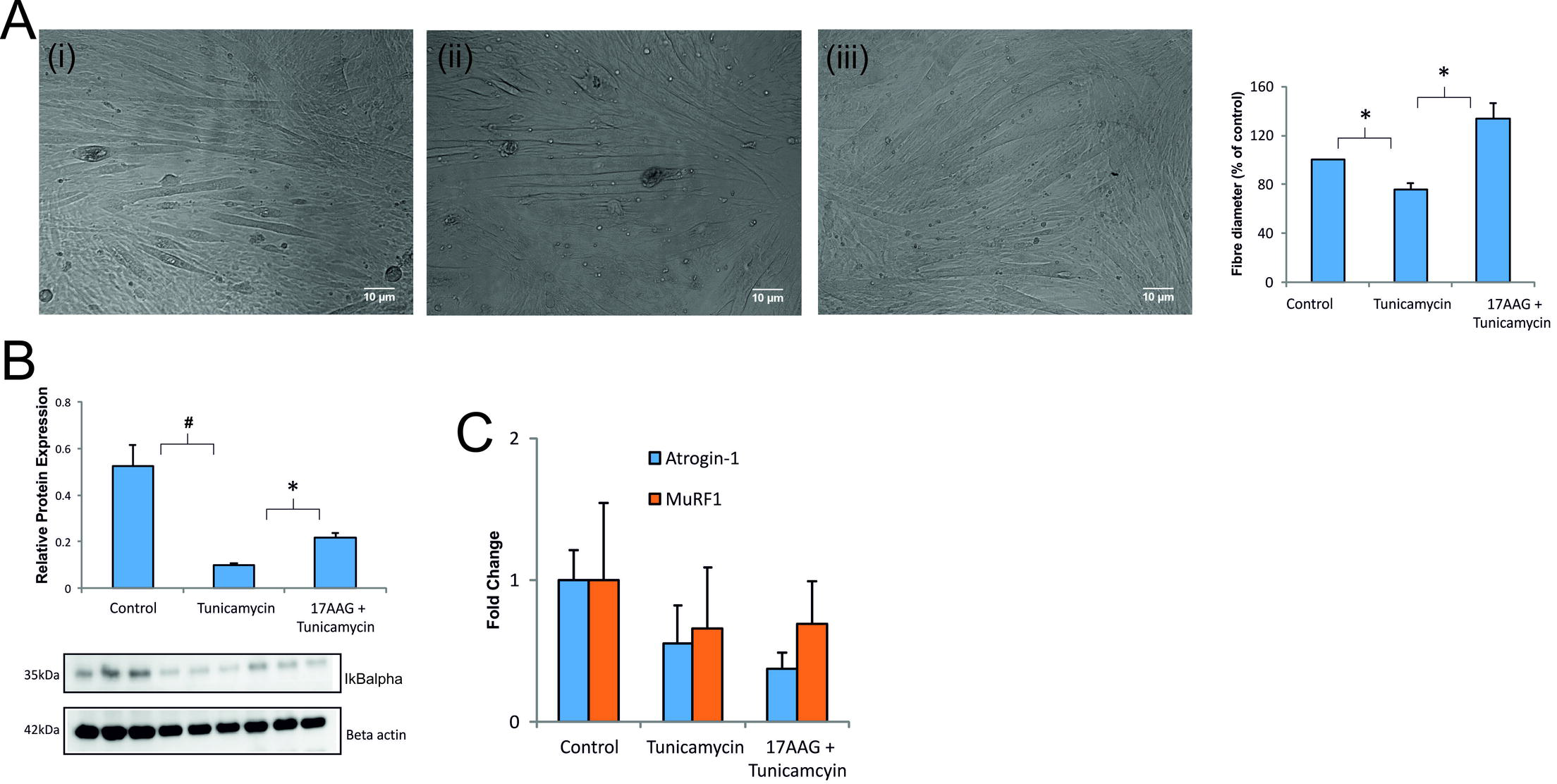
**(A)** Representative light microscopy images (x10 magnification, 10μm scale bar) **(i)** control, **(ii)** Tunicamycin **(iii)** 17AAG + Tunicamcyin, and quantified myotubular diameters – Tunicamycin induced significant fibre shrinkage, i.e. atrophy, an effect minimised by simultaneous 17AAG treatment; **(B)** Quantified densitometry and representative western blot images of IκBα – Tunicamycin induced obvious reductions in IκBα protein content, an effect prevented by 17AAG treatment; **(C)** qPCR data showing gene expression levels of *Atrogin-1* and *MuRF-1*, which showed no significant changes in response to Tunicamycin. Data presented are mean ± SEM (n=3-6) *p≤0.05 ^#^p≤0.01.

### 17AAG protects against ER stress-induced mitochondrial dysfunction, and is associated with JNK phosphorylation

Based on the knowledge that ER stress-induced mitochondrial dysfunction is associated with JNK phosphorylation, we investigated whether HSP70 up-regulation could modify this interaction, [12]. Tunicamycin treatment significantly impaired mitochondrial function, as evidenced by reductions in the RCR and P:O ratios, and in total ATP levels, effects all ameliorated by 17AAG (**Figures 4A-C**). However, and perhaps surprisingly, we observed no changes in mitochondrial electron transport chain (ETC) complex density (**Figure 4D**), nor in any of the genes associated with mitochondrial biogenesis, fusion or fission (**Figures 4E**). At the same time, phosphorylated levels of JNK were increased with Tunicamycin treatment, an effect lessened by 17AAG (**Figure 4F-G**).

**Figure 4:**
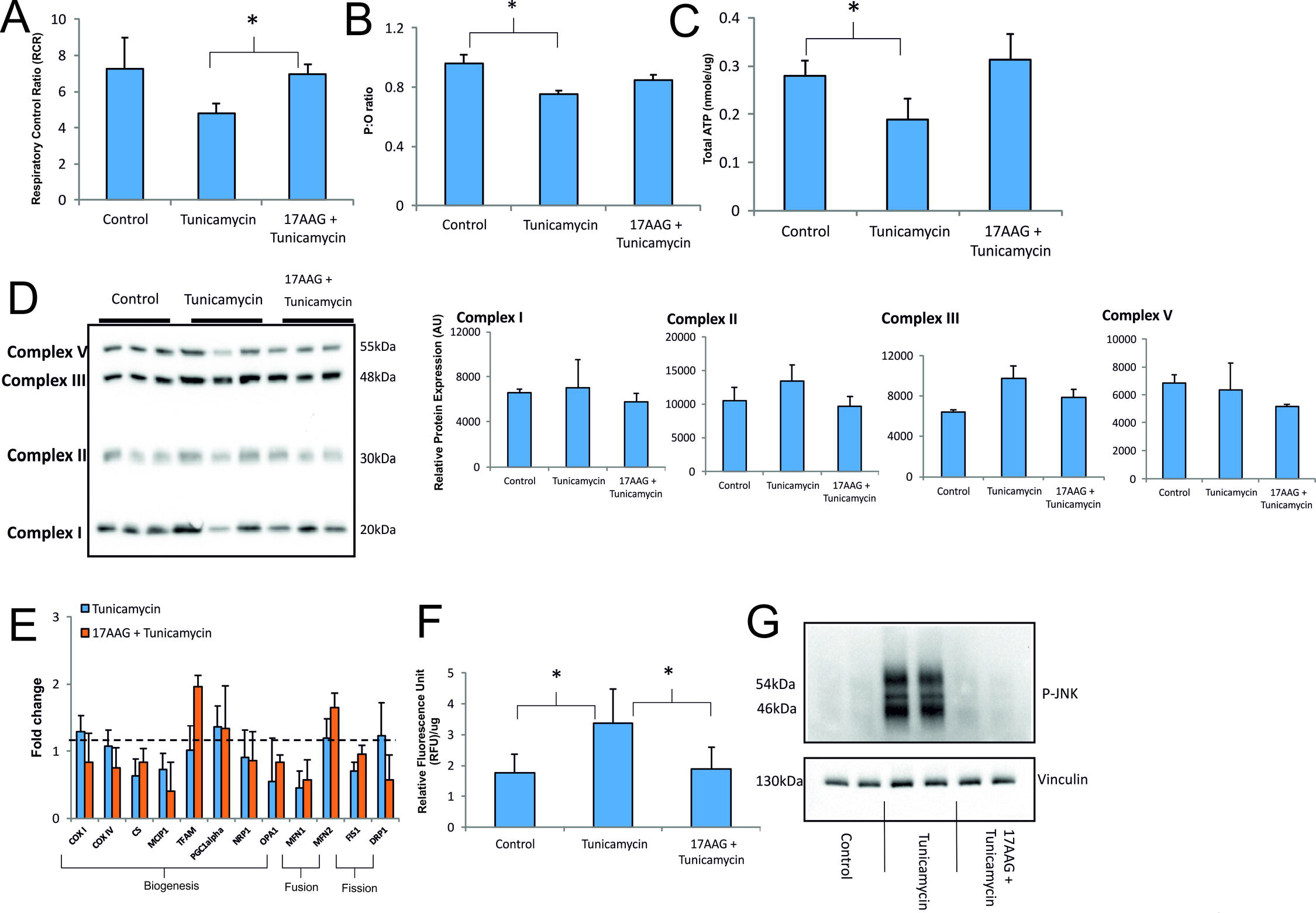
**(A)** Tunicamycin partially reduced the respiratory control ratio (RCR) and **(B)** significantly reduced the phosphate:oxygen (P:O) ratio and **(C)** the total ATP levels, these affects ameliorated by 17AAG. **(D)** Mitochondrial complex density and representative western blot images showed no significant Tunicamycin-induced changes and **(E)** Shows expression levels of genes associated with the mitochondrial dynamics of biogenesis, fusion and fission: i.e. *MFN1*, *MFN2*, *COX I*, *COXIV*, citrate synthase, *MCIP1*, *TFAM*, *PGC1-alpha*, *NRP1*, *OPA1*, *FIS1* and *DRP1* (See list of abbreviations), [23]. Genes associated with dynamics were all unaltered by Tunicamycin; **(F)** levels of phosphorylated JNK detected by ELISA; **(G)** representative western blot of phosphorylated JNK – Tunicamycin induced elevation in P-JNK, an effect prevented by 17AAG treatment. Data presented are mean ± SEM (n=3-6) *p≤0.05.

### ER stress-induced mitochondrial dysfunction is associated with increased markers of oxidative stress and ETC uncoupling

Gene expression levels of *UCP-3* were elevated following Tunicamycin treatment, suggesting that the ETC had become uncoupled in response to ER stress. This effect was attenuated by 17AAG (**Figure 5A**). Total thiol levels were reduced by Tunicamycin treatment, clearly suggesting that increased ROS generation had occurred in response to ER stress, an effect significantly reduced by 17AAG (**Figure 5B**). Similarly, H_2_O_2_ levels were elevated in response to ER stress and attenuated in the presence of 17AAG (**Figure 5C**). Gene expression levels of the antioxidant enzymes *SOD2* and *PDRX3* were significantly increased by Tunicamycin treatment, again suggesting an elevated generation of ROS in response to ER stress, with a significant decline in *PDRX3* but not *SOD2* gene expression in the presence of 17AAG (**Figures 5C-D**).

**Figure 5:**
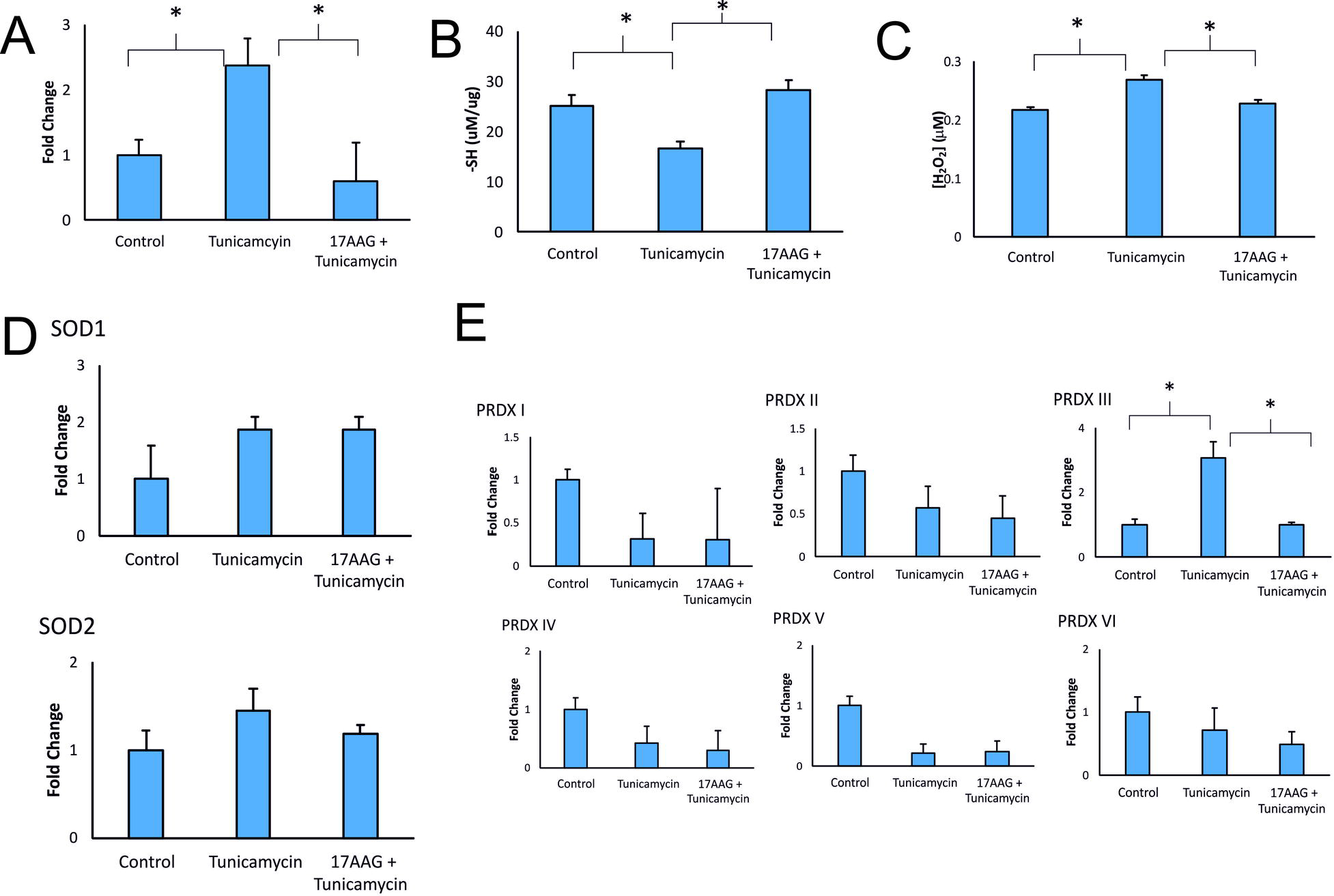
**(A)** Gene expression levels of *UCP-3* were significantly increased by Tunicamycin, an effected blocked by 17AAG treatment, as were **(B)** Total thiol levels, **(C)** H_2_O_2_ levels. Gene expression levels of **(D)** *SOD1* and *SOD2* and **(E)** of *PRDX I-VI* were increased by Tunicamycin treatment, an effect blocked by 17AAG treatment. Data presented are mean ± SEM (n=4-6) *p≤0.05.

## Discussion

The ER stress pathway is an important component of the normal cellular protein folding machinery, permitting cells to manage perturbations in protein homeostasis. However, when chronically over-activated, the ER stress pathway also plays an important role in mediating various pathologies, [24]. Patients with myositis exhibit chronic muscle ER stress pathway over-activation and this appears likely to play a pathogenic role, although the precise mechanisms remain to be elucidated. Given the mutual proximity of the ER and mitochondria in muscle cells, and the capability of the ER stress pathway to modify mitochondrial bioenergetics and induce ROS generation, others and we have suggested that these factors are a feasible cause of non-immune cell-mediated weakness induction in myositis, [3, 8]. Based on the cytoprotective properties of 17AAG in a range of myopathologies other than myositis, we speculated that this agent might reduce ER stress-induced mitochondrial dysfunction and limit potentially harmful ROS generation in skeletal muscle cells.

The *in vitro* model used here appears to mimic components of ER stress pathway activation in myositic muscle, illustrated by observations of elevated protein levels of Grp78 and Grp94 in both *in vivo* human and murine myositis [9], and as illustrated by the increased *XBP-1* gene splicing, following Tunicamycin treatment of murine myotubes, [9, 26]. In the absence of ER stress-induced apoptosis, the latter confirmed by the absence of caspase-12 cleavage, these current results suggest that any bioenergetic changes induced in the presented murine model were not associated with muscle cell death.

Myofibre atrophy is a key pathological feature of myositis. In the model being presented here, ER stress induced significant myotubular atrophy; but this was preventable by the pharmacological deployment of 17AAG. In view of the observation of induced myotubular atrophy, we investigated key players thought likely to be involved in atrophy induction, i.e. the atrogenes *Atrogin-1* and *MuRF-1*, and NF-κB activation. We have previously demonstrated the use of IkappaBalpha (IκBα) degradation as a sensitive way of indirectly assessing nuclear factor kappa B (NF-κB) activity in C2C12 myotubes, [27]. Surprisingly, in the current experiments, we observed no Tunicamycin-induced changes in *Atrogin-1* or *MuRF-1* gene expression levels. However, Tunicamycin-induced ER stress did cause significant IκBα degradation, thus confirming that NF-κB activation had occurred, an effect significantly reduced by 17AAG. A role for NF-κB activation in inducing myotubular atrophy is well established in a rodent model of muscle wasting, [28]. Based on these observations we propose that the *in vitro* murine model presented here mimics at least some features exhibited by muscle cells from human myositis patients.

The current results confirm that ER stress pathway activation modifies myotubular oxygen utilisation, as evidenced by declines in the RCR and P:O ratios and falls in total cellular ATP levels. 17AAG prevented or ameliorated these induced bioenergetic deficits. Mechanistically these alterations in mitochondrial function have previously been attributed to phosphorylation of JNK by the ER stress receptor, inositol-requiring enzyme 1 (IRE1) α, with subsequent migration of the P-JNK complex to the mitochondrial membrane, thus causing a depression of the ETC, [12, 29]. ER stress induced ETC depression has also been associated with elevated mitochondrial ROS generation, with its potential to cause oxidative damage to contractile proteins, [10]. In our *in vitro* model Tunicamycin-induced ER stress pathway activation was associated with increased phosphorylation of JNK, an effect clearly reduced by 17AAG. These findings support previous work linking ER stress activation to mitochondrial dysfunction via JNK phosphorylation, [12]. The precise mechanism for the downregulation of JNK phosphorylation is unclear. The activity of 17AAG orchestrated through its inhibition of HSP90 suggests a wealth of client proteins which could mediate these changes [30]. Some evidence has shown that heat shock protein 70 (a HSP90 client protein) can regulate JNK phosphorylation, and this may be a mediator in our model – however, further investigation is needed in this context [31, 32].

Tunicamycin-induced ER stress pathway activation induced no changes in mitochondrial ETC complex density or mitochondrial dynamics genes. These findings suggest that the observed mitochondrial dysfunction was not due to changes in mitochondrial numbers or dynamics. In keeping with the notion that ROS plays some role in mediating the downstream effects of ER stress activation, we observed a decline in the total thiol (sulphydryl) pool and elevations in H_2_O_2_ levels, in combination with induced elevations of gene expression levels of *SOD2* and *PRDX3*. ROS generation in response to ER stress is well characterised, and its role in impairing mitochondrial function in muscle may in part be associated with the activation of uncoupling proteins. In *in vitro* model presented here, we observed elevated *UCP-3* gene expression in response to ER stress. Uncoupling proteins are redox sensitive, thus permitting proton leak across the inner mitochondrial membrane in response to ROS generation, a process uncoupling the link between oxidative phosphorylation and ATP synthesis, [33]. We therefore suggest that ER stress induces P-JNK/ROS-mediated uncoupling of the ETC, causing decreased oxygen utilisation and ATP synthesis, effects clearly ameliorated by 17AAG. Our narrative of ROS involvement in these processes is supported by recent findings in a murine model of myositis, whereby weakness is associated with interferon-γ-induced ROS generation, [34].

## Conclusions

ER stress is thought to play an important role in inducing non-immune cell-mediated muscle contractile and energy deficits, and which are likely to apply in human myositis, [8, 9]. However, the precise mechanisms mediating weakness-induction downstream of ER stress remain unexplained. In the current experiments, we have modelled ER stress *in vitro*, and demonstrated obvious declines in mitochondrial function, but which are mitigated by 17AAG. These observations suggest that chronic ER stress in myositis may induce perturbations in mitochondrial function capable of causing bioenergetic deficits, which likely contribute to muscle weakness-induction.

While recognising the limitations of the use of an *in vitro* model, the link between ER and mitochondria is conserved across multiple species and cell types, so likely also applies in human skeletal muscle cells. Given the crucial role that muscle ER plays in controlling the calcium fluxes required to control complex muscle contractions, as well as controlling energy utilisation, it appears likely that the mechanistic insights gained here do have some relevance for understanding human myositis, and other ER stress-associated myopathologies.

**Supplementary Table 1:** Primer sequences for the genes of interest in the qPCR analyses.

## List of abbrevations

**17AAG** - 17-*N*-allylamino-17-demethoxygeldanamycin; **MCIP1** - Modulatory calcineurin interacting protein 1; **TFAM** - Mitochondrial transcription factor A; **DRP1** – Dynamin-related protein 1; **PGC1α** - Peroxisome proliferator-activated receptor gamma coactivator 1-alpha; **MFN1** – Mitofusin 1; **MFN2** – Mitofusin 2; **COX I** – Cytochrome *c* oxidase subunit I; **COX IV** – Cytochrome *c* oxidase subunit IV; **NRF1** – Nuclear respiratory factor 1; **OPA1** – Optic atrophy type 1; **FIS1** – Mitochondrial fission 1 protein; **Grp78** – Glucose regulate protein 78; **Grp94** – Glucose regulate protein 94 **PRDX** – Peroxiredoxin; **UCP-3** – Uncoupling protein 3; **SOD1** – Superoxide dismutase 1; **SOD2** – Superoxide dismutase 2; **IκBα** – I kappa B alpha; **JNK** - c-Jun N-terminal kinase; **RCR** – Respiratory control ratio; **P:O** – Phosphate: Oxygen; **XBP-1** – X-box binding protein 1

